# A polymerisation-associated conformational switch in FtsZ that enables treadmilling

**DOI:** 10.1101/093708

**Authors:** James M. Wagstaff, Matthew Tsim, María A. Oliva, Alba García-Sanchez, Danguole Kureisaite-Ciziene, José Manuel Andreu, Jan Löwe

## Abstract

Bacterial cell division in many organisms involves a constricting cytokinetic ring that is orchestrated by the tubulin-like protein FtsZ. FtsZ forms dynamic filaments close to the membrane at the site of division that have recently been shown to treadmill around the division ring, guiding septal wall synthesis.

Here, using X-ray crystallography of *Staphylococcus aureus* SaFtsZ we reveal how an FtsZ can adopt two functionally distinct conformations, open and closed. The open form is found in SaFtsZ filaments formed in crystals and also in soluble filaments of *E. coli* FtsZ as deduced by cryoEM. The closed form is found within several crystal forms of two non-polymerising SaFtsZ mutants and corresponds to many previous FtsZ structures from other organisms.

We argue that FtsZ undergoes a polymerisation-associated conformational switch. We show that such a switch provides explanations for both how treadmilling may occur within a single-stranded filament, and why filament assembly is cooperative.

## Introduction

FtsZ is an ancient, filament forming, tubulin-like GTPase protein found in the vast majority of bacteria and archaea, where it acts as a central component of the cell division machinery (1–3). FtsZ is localised to the plasma membrane at future division sites resulting in the emergence of a ring structure around the centre of the cell, the Z-ring. FtsZ is anchored to the plasma membrane by other proteins, most often FtsA but also ZipA and/or SepF (4–6). FtsA is a divergent actin homologue that forms copolymers with FtsZ and contains an amphipathic helix that facilitates membrane attachment (7).

After the localisation of FtsZ, a large number of other proteins are recruited to the division site. These proteins carry out remodelling and synthesis of cell wall during the division process. Together these proteins have been termed the divisome, although it is currently not clear which components, if any, form stable multi-subunit complexes during the process.

The precise molecular architecture of the Z-ring remains unclear, although it is probably composed of dynamic overlapping filaments along the circumference of the ring, at least during the later stages of the division process in rod-shaped model organisms such as *Escherichia coli* (8). It was already clear from early fluorescence microscopy studies that during the cell division process the Z-ring contracts with the invaginating septum (9). *In vitro* reconstitution experiments of FtsZ and FtsA with membranes showed that these two components alone deform membranes (10). Together with homology to force-generating eukaryotic tubulins this prompted the suggestion that FtsZ has a role in generating forces required for invagination. In contrast, observations of constrictions and divisions of cells with helical Z-rings, incomplete Z-rings, and divisomes with modified FtsZ properties, support the opposing idea that FtsZ does not provide an indispensable driving force for constriction (11–14). The alternative candidate for force generation is cell wall remodelling. A third option is that cell wall remodelling and Z-ring dynamics are interlinked processes that work together to generate the forces needed for division to occur robustly and efficiently every time.

Recently, *in vitro* treadmilling of FtsZ filaments has been reported on supported bilayers (15, 16) and also *in vivo* where FtsZ filaments were found to treadmill with components of the divisome around the division site (17, 18). These findings have resurrected an old model of bacterial cell division: the template model, in which the closing septum constricts by new cell wall material being deposited in concentric rings on the inside of old material by moving synthesis machinery, which in turn is guided or organised into a ring by dynamic FtsZ filaments. This idea fits into the third category of ideas about the role of FtsZ.

Treadmilling is a property of certain cytomotive filaments whose filament-forming subunit interfaces are made dynamic in time through nucleotide hydrolysis that is, in turn, triggered by the polymerisation reaction. Treadmilling requires a difference in the rate of net polymerisation and de-polymerisation at the so-called plus and minus ends of the filaments. FtsZ so far was not considered a good candidate for treadmilling behaviour largely because it is currently not known if any well ordered structures are formed in cells beyond single-stranded protofilaments. Differing rates of subunit addition at the two ends of simple single-stranded filaments are difficult to envisage (19).

Surprisingly, knowledge of FtsZ filament architecture is limited. Only one FtsZ crystal form, from *Staphylococcus aureus* (SaFtsZ, PDB IDs 3VOA, 3VO8), has revealed a straight protofilament of FtsZ, as might be expected from electron micrographs of many different FtsZ filaments (20). The arrangement of this crystalline filament is very similar to protofilaments formed by tubulin, the eukaryotic cousin of FtsZ. Tubulin protofilaments are completely straight in microtubules, and have special sequence regions, such as the M-loop (missing in FtsZs), which make lateral interactions between protofilaments. The conformation of SaFtsZ subunits in those straight filaments showed an unusually (as compared to other FtsZ structures) open conformation, with the N-terminal GTP binding domain (NTD) and C-terminal GTPase activation domain (CTD) being rotated and shifted apart (∼27°). Subsequent crystallisation efforts using constructs with large changes to the critical T7 loop that normally contacts the GTP/GDP nucleotide bound to the next subunit were successful in generating crystals where SaFtsZ adopted a different ‘closed’ conformation, more similar to FtsZs from other species (PDB IDs 3WGK, 3WGL) (21). These crystals contained straight protofilaments. However, it remained unclear what caused the switch and whether it was dominated by non-specific crystal contacts or the alterations of the T7 loop.

Two of the previous open SaFtsZ structures also include the FtsZ functional inhibitor and filament stabiliser PC190273 (PDB IDs 3VOB, 4DXD), bound in the cleft between the N-and C-terminal domains, only possible in the open conformation - presumably locking the protein in this state (20, 22). PC190273 reduces polymerisation cooperativity *in vitro*(23). Isolated open form SaFtsZ monomers relax into the closed conformation during molecular dynamics simulations (24). Fluorescent analogues of PC190723 have recently been used to monitor opening and closing of the inter-domain cleft in solution as a function of FtsZ polymerisation state (25). Together, these results hint that the closed form of FtsZ seen in many crystals is the predominant conformation of monomeric FtsZ and *vice versa* that filamentous FtsZ in solution is in the open conformation seen in SaFtsZ filament crystals. Currently what is lacking is robust structural evidence that this is the case.

FtsZ shares two properties with actin and tubulin that until now have been hard to explain. Firstly, FtsZ exhibits cooperative assembly, with a critical concentration and a lag phase in assembly. This is not possible for a single-stranded, isodesmic filament with rigid subunits, and an assembly switch has long been hypothesised to explain this cooperativity (26). Secondly, filament treadmilling is presumed to require multi-strandedness, which FtsZ may not have.

Here we demonstrate that FtsZs do indeed undergo a conformational switch, that this switch is associated with polymerisation and not nucleotide hydrolysis state and that switching provides a possible mechanism for both cooperative assembly and also for treadmilling, which has been proposed to be a key dynamic filament behaviour used to organise cell wall remodelling.

## Results and Discussion

### SaFtsZ-T66W and −F138A are polymerisation and GTPase compromised

Two SaFtsZ mutations, F138A and T66W were designed to inhibit SaFtsZ filament formation, based on equivalent mutations inhibiting assembly of *Methanocaldococcus jannaschii* FtsZ (M164A (27) and T92W (28) respectively, polymerisation inhibition of T92W unpublished data). Both mutation sites are located on the ‘top’ surface of FtsZ, on the N-terminal, GTP-binding domain and are part of the longitudinal protofilament interface seen in crystals. Full-length, untagged, SaFtsZ wildtype, F138A, and T66W proteins were purified and characterised biochemically (Figure 1). Filament formation in both SaFtsZ mutated proteins was compromised since no filament formation was detected by sedimentation (Figure 1A) or negative stain electron microscopy (Figure 1C) for either T66W or F138A in the presence of GTP or guanosine-5’-[(α,β)-methyleno]triphosphate (GMPCPP), a slowly-hydrolysable analogue of GTP. FtsZ GTPase activity is largely dependent on polymerisation as one subunit provides catalytic residues to the active site of the next subunit through residues in loop T7. Both mutants have weak GTPase activity (Figure 1B), indicating that monomers may at least associate to form transient but functional active sites. In support of this, on addition of PC190723, the mutant proteins did form filaments detectable by sedimentation and electron microscopy in the presence of GTP and GMPCPP. We conclude that SaFtsZ T66W and F138A are polymerisation and GTPase compromised but retain some residual activities.

**Figure 1.**
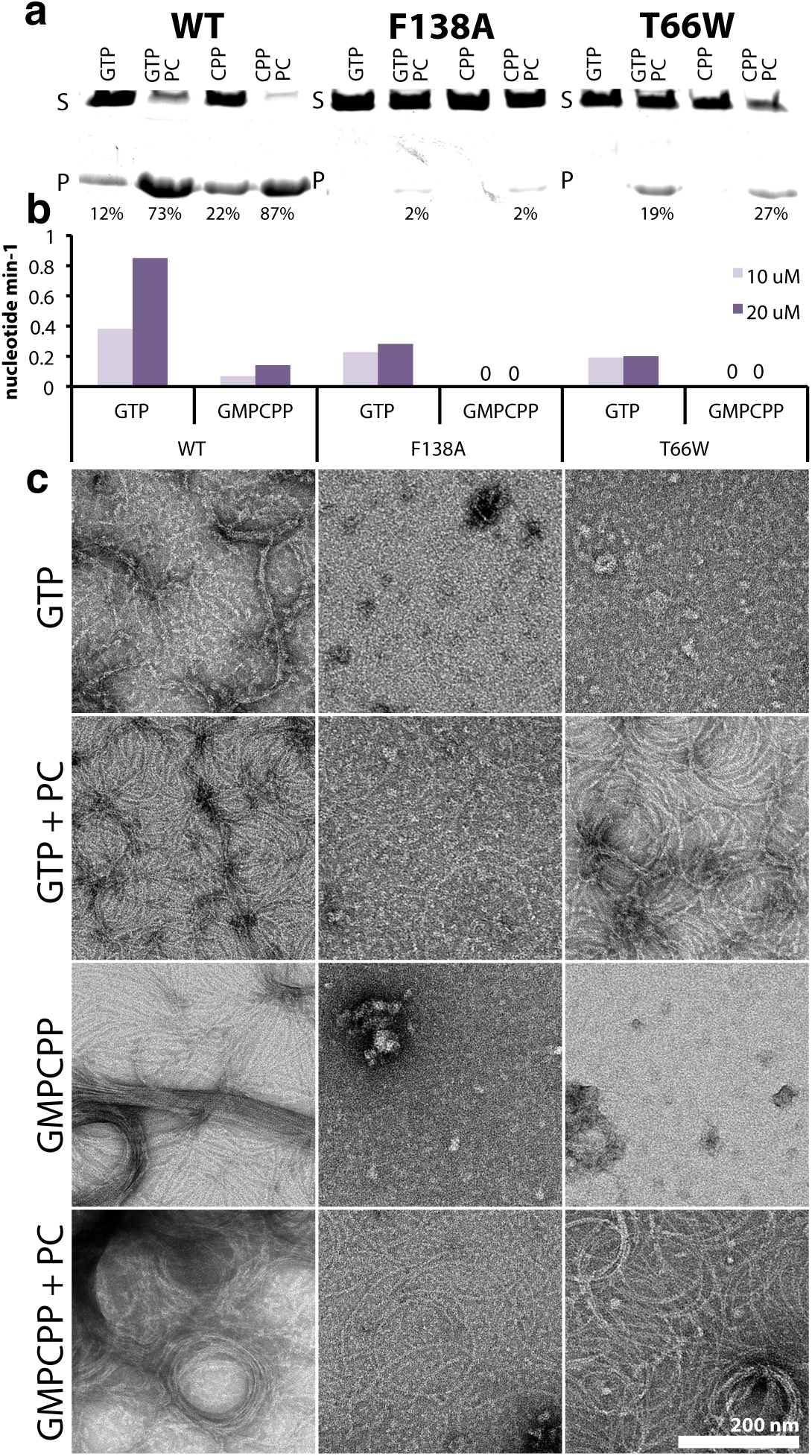
SaFtsZ full-length mutant proteins T66W and F138A have compromised polymerisation and GTPase activities. **A)** Polymerisation of FtsZ proteins at 10 μM was assayed by sedimentation in the presence of GTP and GMPCPP (CPP) with and without FtsZ inhibitor PC190723 (PC). Pelleted (P) and soluble (S) protein was run in the same lane of an SDS-PAGE gel with a delay. Percentage of pelleted protein was estimated from integration of band intensities. **B)** GTPase activity of FtsZs at 10 and 20 μM in the presence of GTP/GMPCPP. **C)** Polymerisation of FtsZ proteins in the presence of GTP and GMPCPP with and without FtsZ inhibitor PC190723 (PC) was assessed by negative stain electron microscopy. All images are at the same magnification (scale bar 200 nm).

### SaFtsZ adopts either a closed or an open conformation

We solved five crystal structures of the globular domains of SaFtsZ T66W and F138A (Table S1, Figure 2). SaFtsZ constructs truncated to residues 12-316 were used to remove the N and C-terminal tails of FtsZ previously found to inhibit crystallisation. For easier reference, the five SaFtsZ structures are named herein in the form #XXx: number (1–5), mutation (F for F138A, T for T66W), monomer conformation (O for open, C for closed), and finally the arrangement of monomers within the crystal (m for monomeric, f for filamentous, single protofilament and s for split/domain swapped).

**Figure 2.**
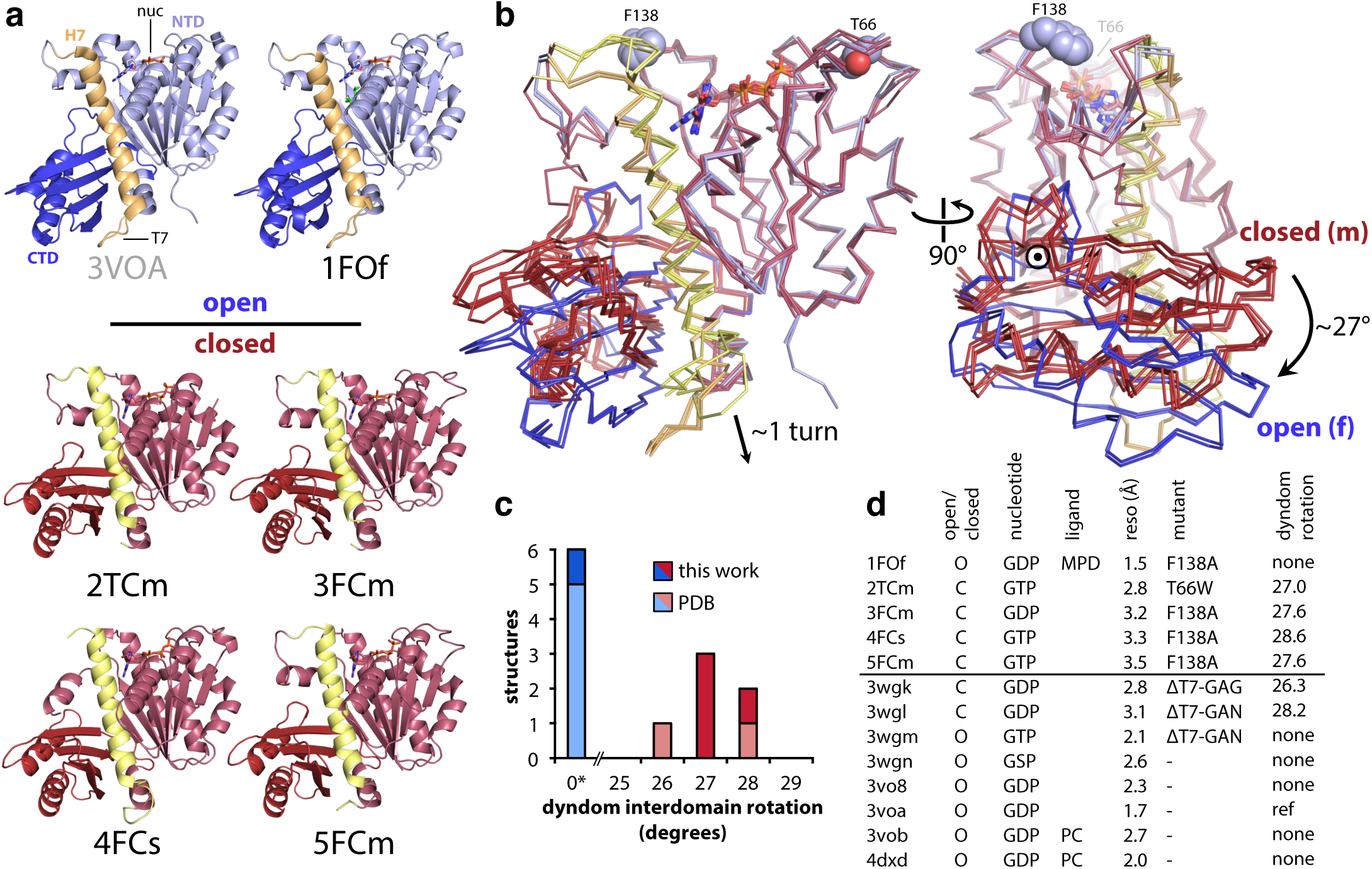
Nucleotide-bound SaFtsZ crystal structures grouped into two conformations: open and closed. **A)** The five SaFtsZ (truncated to residues 12-316) structures determined here and PDB ID 3VOA are shown in cartoon representation, with nucleotides as sticks coloured by element. MPD molecule in 1FOf is shown as green sticks. The structures are coloured according to conformation. Closed structures are shown in red, with the N−terminal GTP-binding domain in light red, and the Cterminal GTPase activation domain in dark red, central helix H7 is highlighted in yellow. Open structures are shown in blue, with the N−terminal domain in light, and the C-terminal domain in dark blue, central helix H7 is highlighted in orange. All structures shown in the same orientation, after alignment to the N−terminal domain of 3VOA (residues 13-165). 4FCs domain-swapped pseudomonomer is formed of two polypeptides. Note the different position of the C-terminal domain in the two sets of structures. **B)** Superposition of the six structures in (A) shown in Cα ribbon representation, after alignment as for (A), with the same colour scheme. Nucleotides are shown as sticks. Sidechains of residues F138 and T66 of wildtype structure are shown as spheres, non-carbon atoms coloured by element. (left) the same view as in (A), (right) molecules rotated 90° as indicated. Axis of interdomain rotation indicated by ‘¤’, direction by arrow. **C, D) Census of available nucleotide-bound SaFtsZ structures. C)** Bar chart indicating that DynDom, model-free assessment of dynamic protein domains, essentially produces two groups when comparing SaFtsZ structures to PDB ID 3VOA: no interdomain rotation, or a ∼27° shift (around the axis in (B, right). **D)** Table with information about nucleotide-bound SaFtsZ structures. Horizontal line separates structures determined here (above) from previous structures in the PDB. (MPD: 2-methyl-2,4-pentanediol, PC:
PC190723).

**1FOf** (PDB ID 5MN4) is a structure of SaFtsZ F138A at a resolution of 1.5 Å, with one molecule of FtsZ in the asymmetric unit (ASU), a GDP molecule in the active site and also a molecule of 2-methyl-2,4-pentanediol (MPD) which was present at high concentration in the crystallisation drop. Residues 12-315 are present in the model. **2TCm** (PDB ID 5MN5) is a structure of T66W at a resolution of 2.8 Å, there are two molecules of FtsZ in the ASU with a GTP molecule in the active site of both molecules. Residues 13-62, and 74-315 are present in chain A of the model, the missing residues are in the T3 loop, part of the nucleotide-binding pocket (including the mutated residue). The same residues are seen in chain B, except 203-208, in the T7 loop, are also missing. **3FCm** (PDB ID 5MN6) is a structure of F138A at a resolution of 3.2 Å, there are two molecules of FtsZ in the ASU with a GDP molecule in the active site of both molecules. Residues 13-201, and 209-315 are present in chain A of the model, the missing residues are in the T7 loop. Residues 12-315 are present in chain B of the model. **4FCs** (PDB ID 5MN7) is a structure of F138A at a resolution of 3.3 Å, there are two molecules of FtsZ present in the ASU although the N−and C-terminal domains of each polypeptide have become disengaged and reformed in pseudo-complete FtsZ molecules with the corresponding domains of crystallographic-symmetry related molecules, in other words a domain swap. Residues 12-315 are present in both chains in the model. There is one molecule of GTP for each chain, and the position of the coordinating magnesium ion was identified for each. **5FCm** (PDB ID 5MN8) is a structure of SaFtsZ F138A at a resolution of 3.5 Å, there are two molecules of FtsZ in the ASU with a GTP in the active site of both molecules. Residues 13-202, and 207-315 are present in both chains of the model, the missing residues are in the T7 loop.

One structure (1FOf) was in the open form and essentially identical (crystallographically isomorphous) to previously published wildtype SaFtsZ open conformation structures (Cα RMSD versus PDB 3VOA: 0.33 Å) (Figure 2A, top). FtsZ molecules in 1FOf form completely straight single-stranded filaments (protofilaments) with a 44 Å repeat extending throughout the crystal. Four of the polymerisation compromised FtsZ point mutant structures (2FCm, 3FCm, 4FCs, 5FCm) were in the alternative closed conformation previously seen in SaFtsZ after extensive mutation of the T7 loop (e.g. Cα RMSD 2TCm vs PDB ID 3WGL: 1.50 Å). Indeed, the closed structures were successfully solved by molecular replacement with one of the previous T7 loop replacement mutant structures (PDB 3WGL) as the starting search model. Unlike the closed form T7 mutant SaFtsZ crystals, none of our closed crystal forms contain straight filaments running through the crystals.

When we analysed the conformations of all of the available nucleotide-bound SaFtsZ structures it became clear that they do fall into two discrete groups (Figure 2B-D). We excluded SaFtsZ apo structures (e.g. PDB ID 3VO9), which are very different and are unlikely to be physiologically relevant given the high concentration of GNP in cells. The two nucleotide-bound conformations can be distinguished clearly by the change of the interdomain angle between the N−and C-terminal domains. If we consider the NTD to be fixed in space, the switch to the open conformation is best defined (as determined by the model-free program DynDom (29)) as a ∼27° rotation of the CTD versus the closed conformation, around an axis of rotation as indicated (⊙) in Figure 2B, right. This rotation is accompanied by a downward shift of the central helix 7 (H7, yellow in Figure 2) by almost one helical turn (Figure 2B, left).

It is important to note that, including previous work, we have now available SaFtsZ structures with all permutations of open/closed conformations and bound GTP/GDP nucleotide. Also, for the first time, with this work we have structures showing an identical near-native FtsZ molecule in multiple conformations: SaFtsZ F138A crystallised in the open conformation in straight filaments as 1FOf, in the closed form as a monomer in two different space groups in 3FCm and 5FCm and as a split/domain-swapped closed form monomer in 4FCs. Domain swapping has been seen before for FtsZ (30), and highlights the surprising independence of the N−and C-terminal domains.

Based on previous work that established that all FtsZ structures are broadly similar (31) we decided to compare all FtsZ structures to all others, including the ones presented here. The striking similarity of all FtsZ structures, except SaFtsZ open forms, is illustrated in Figure 3. There are many interesting results in Figure 3C, but the most significant is the fact that SaFtsZ closed forms have a more similar conformation to other FtsZs, even archaeal ones, than to SaFtsZ open structures. The most obvious outlier to the overall trend is PDB ID 1W5F, the previously published structure of a domain-swapped FtsZ from the extremophile bacterium *Thermotoga martima*. In the alignment in Figure 3A and B the 1W5F structure can be easily identified as it falls approximately between the two clusters. Also, our domain-swapped structure 4FCs aligns relatively poorly to the other closed structures, although is much more similar to closed structures than open ones. Both cases are perhaps unsurprising, as domain swapping will clearly impose additional constraints on the conformational freedom of the protein. In the case of 1W5F the two swapped monomers contact one another via their CTDs, so it is unlikely to represent a *bona fide* intermediate form.

**Figure 3.**
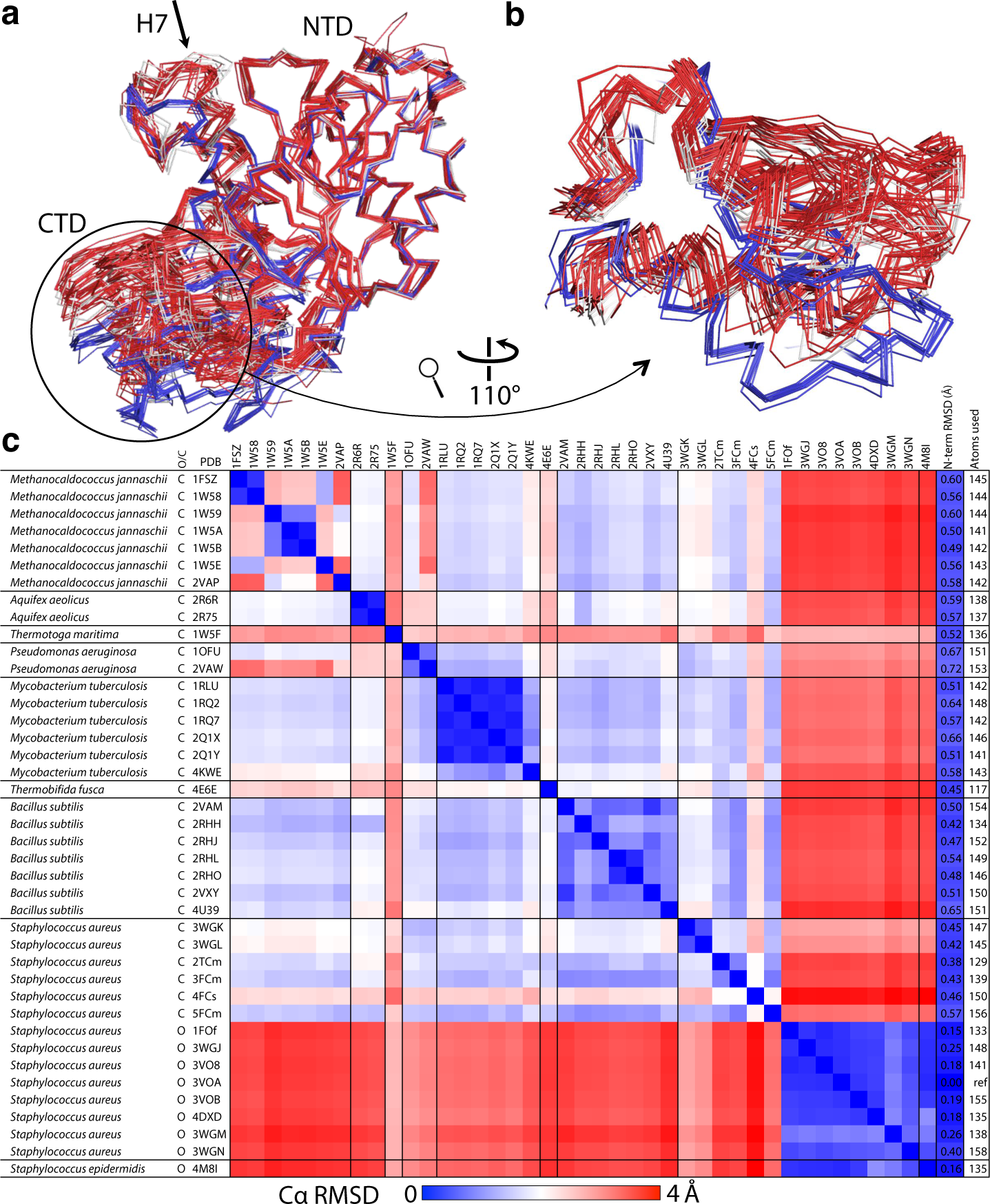
All FtsZ structures (except apo SaFtsZs) can be grouped into two conformations, open and closed. **A)** All previous FtsZ structures were obtained from the PDB as listed in (B). Chain A from each downloaded structure, and the five structures determined here, was extracted and aligned to the N−terminal domain (residues 12 −176) of 3VOA using the PyMOL align command (matches residues via sequence then minimises RMSD (root mean square distance) with 5 cycles of outlier rejection; except for PDB ID 1W5F and our structure 4FCs which are both domain-swapped, in these cases a pseudomonomer was generated for each, also *S. aureus* apo-structures (PDB IDs 3VO9, 3VPA), which have a very different conformation (20), were excluded. N−and C-terminal extensions were removed, and the aligned structures are shown in ribbon representation from the same view as in Figure 2B. Closed structures are coloured red, except for closed *S. aureus* structures which are in white, open structures are coloured blue. The structural conservation of FtsZs is clear from the quality of alignment at the N−terminal domain (the outlier-excluded RMSDs, and the number of Cα used is given in the last two columns of (C)). The two groups of structures can be distinguished because of the relative motion of the C-terminal domain — the open blue structures are separated from the closed white and red ones. **B)** The discrete distinction between the two groups is made clearer by zooming into the C-terminal domain as indicated. **C)** Cα RMSDs were calculated for all structures vs all structures, using the PyMOL align command with 0 cycles of outlier rejection (i.e. all residues matched via sequence are included in RMSD calculation). The RMSD for each pair of structures is indicated with a linear 3 colour gradient as indicated below the matrix. Within each species sets of highly similar structures are found (blue squares on the diagonal filling the black lines), with the exception of *S. aureus* where the two conformations, open and closed, align poorly. The *S. aureus* closed structures are more similar to FtsZs from other species than they are to open *S. aureus* structures, indicating that all existing non *S. aureus* FtsZ structures show a similar, closed, conformations.

We conclude that SaFtsZ exists in two distinct conformations, open and closed, and the closed form is much more similar to all other FtsZ structures than to the SaFtsZ open conformation.

### Crystal structures of polymerisation compromised SaFtsZ mutants reveal structural features of the conformational switch

In order to have two stable globular conformations FtsZ must have structural features that rearrange during switching. These features are best visualised using morph (32) interpolations between the structures shown in Figure 2B, as in Video S1. The large rotation of the C-terminal domain (CTD) versus the N−terminal domain (NTD) requires local rearrangement of residues in order to maintain hydrophobic contacts, side chain solvation state and generally favourable intramolecular interactions. The degree of local rearrangement required is reduced by the movement of H7, which moves to stagger rearrangement across the two domain’s interior faces. Displacement of the C-terminal portion of H7 versus the CTD is facilitated by a large hydrophobic region on the interior face of the CTD’s beta-sheet being able to rotate against hydrophobic residues on H7. Side chain rearrangements here are relatively minor. Regions of greater rearrangement around H7 are highlighted in Figure 4, where structures 1FOf and 5FCm are compared (and measurements refer to this pair) although all of the changes discussed are similar in any of our pairs of open/closed structures. Figure 4A and B show rearrangements at the NTD-facing side of H7, around the nucleotide pocket. Notably, when shifting from closed to open Arg 29 moves from the solvent exposed side of H7 to become slotted between H7 and the NTD in the open state (a 6.5 Å displacement of the guanidinium carbon), interacting directly with both guanosine and Asp 187 (on H7, in the closed state itself interacting with the base). R29-E187 is a conserved ion pair in many FtsZs (27) – here we give insight into why. Despite the downward movement of H7, Phe 183 (also on H7) maintains favourable π-stacking with guanosine, because the base rotates around the C1’-N9 bond. The switch from closed to open leads to disruption of ionic interactions across the C-terminal part of H7 and NTD residues, however a subtle rearrangement takes place to maintain a base-base interaction (33) between Arg 191 and His 33 (Figure 4B, inset): the flexibility and length of the Arg sidechain is used to allow the head group to remain almost fixed despite the ∼4 Å movement of the Cα atom.

**Figure 4.**
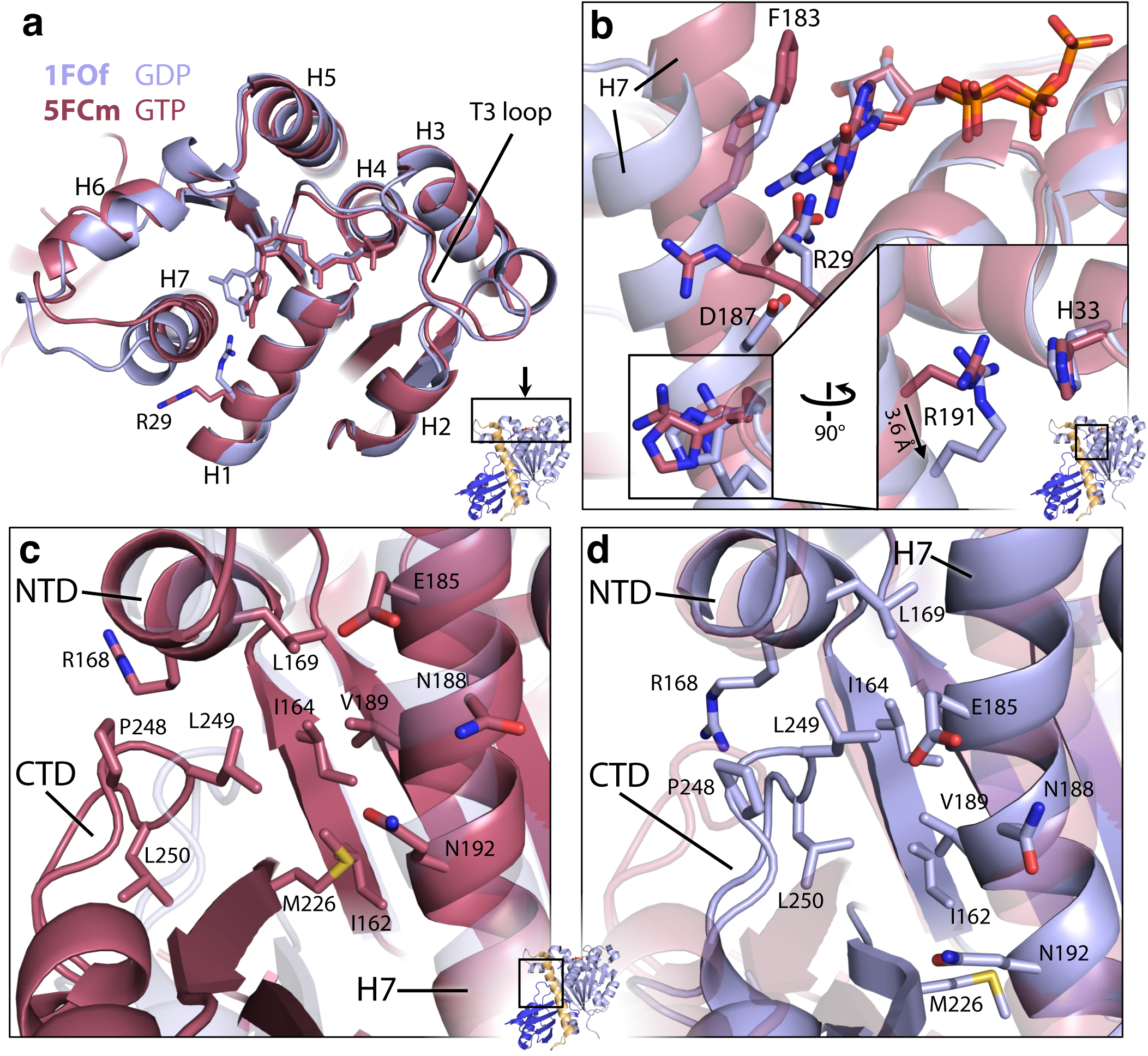
Atomic details of the SaFtsZ regions allowing the conformational switch. Structures 1FOf (open, blue) and 5FCm (closed, red) are shown superposed in cartoon representation after alignment on the N−terminal domain (NTD). Nucleotides and labelled residues are shown as sticks. Non-carbon atoms are coloured by element, except in (A). Viewpoint is indicated in small cartoons, colouring as for Figure 2. **A)** Top view of FtsZ N−terminal domain. Helices are numbered. Note the very minimal rearrangements in this region after both conformational switch and nucleotide hydrolysis. **B)** View of the top of H7 and into the nucleotide binding pocket. Cartoons are semi-transparent. Inset is at same scale and shows molecule 90° rotated as indicated. 3.6 Å is shift of R191 Cα. Note rearrangement of individual sidechains between conformations. **C, D)** Identical views of the three-way interaction between the NTD, C-terminal domain (CTD), and H7, at the top of H7. In (C) 1FOf is semi-transparent with no sidechains, in (D) 5FCm. Identical sidechains are shown in both. The three-way interaction is different in each conformation but solvent is excluded from the hydrophobic core in both cases.

Figure 4C and D illustrate rearrangement in the three-way interactions between the N−terminal part of H7, the NTD, and the CTD. While the residues from the NTD involved in the three-way contact remain relatively fixed in the shift from closed to open, e.g. Leu 169, residues from the CTD beta-sheet and H7 move downward in a coordinated fashion and a loop (246-258) from the CTD loosens allowing residues 248-250 to move towards H7, maintaining solvent exclusion from the hydrophobic pocket.

The conformational switch does not appear to involve structural changes around the phosphate-binding region of the nucleotide-binding pocket (Figure 4A). In particular, the T3 loop can be ordered in all permutations of nucleotide (GDP/GTP or GTPγS) and conformation (open/closed) (1FOf, 3FCm, 5FCm, PDB ID 3WGN), and can even be disordered when GTP-bound (2TCm). These observations appear inconsistent with simple mechanisms of hydrolysis-associated FtsZ monomer conformational change (28, 34), although the terminal phosphate may modulate protein dynamics in a non-obvious way.

### Closed forms of FtsZs correspond to the free monomer, and open forms to the polymerised subunit

We have classified our five polymerisation-compromised mutant SaFtsZ structures as having closed or open conformations, but they can also be grouped according to how they are found in relation to other molecules in the crystal. SaFtsZs in 1FOf, and in previously published open forms (PDB IDs 3WGM, 3VOA, 3VOB, 3VO8, 4DXD) are arranged in straight, single, tubulin-like filaments extending through the crystal – indicated in our naming scheme by the second, lower-case, ‘f’ in 1FOf. Adjacent subunits from each of 1FOf and PDB ID 3VOA, extracted from the crystal lattices (constructed using crystallographic symmetry operators), are shown in Figure 5A. Like in tubulin, the nucleotide forms a large part of the interface between subunits, and it is thought that nucleotide hydrolysis is used to modulate interface affinity. These crystalline filaments have a 44 Å repeat, which corresponds well to repeat intervals seen in negatively stained FtsZ filaments from a number of species. As a result, it has been hypothesised that the 1FOf-like crystal filaments resemble soluble FtsZ filaments.

**Figure 5.**
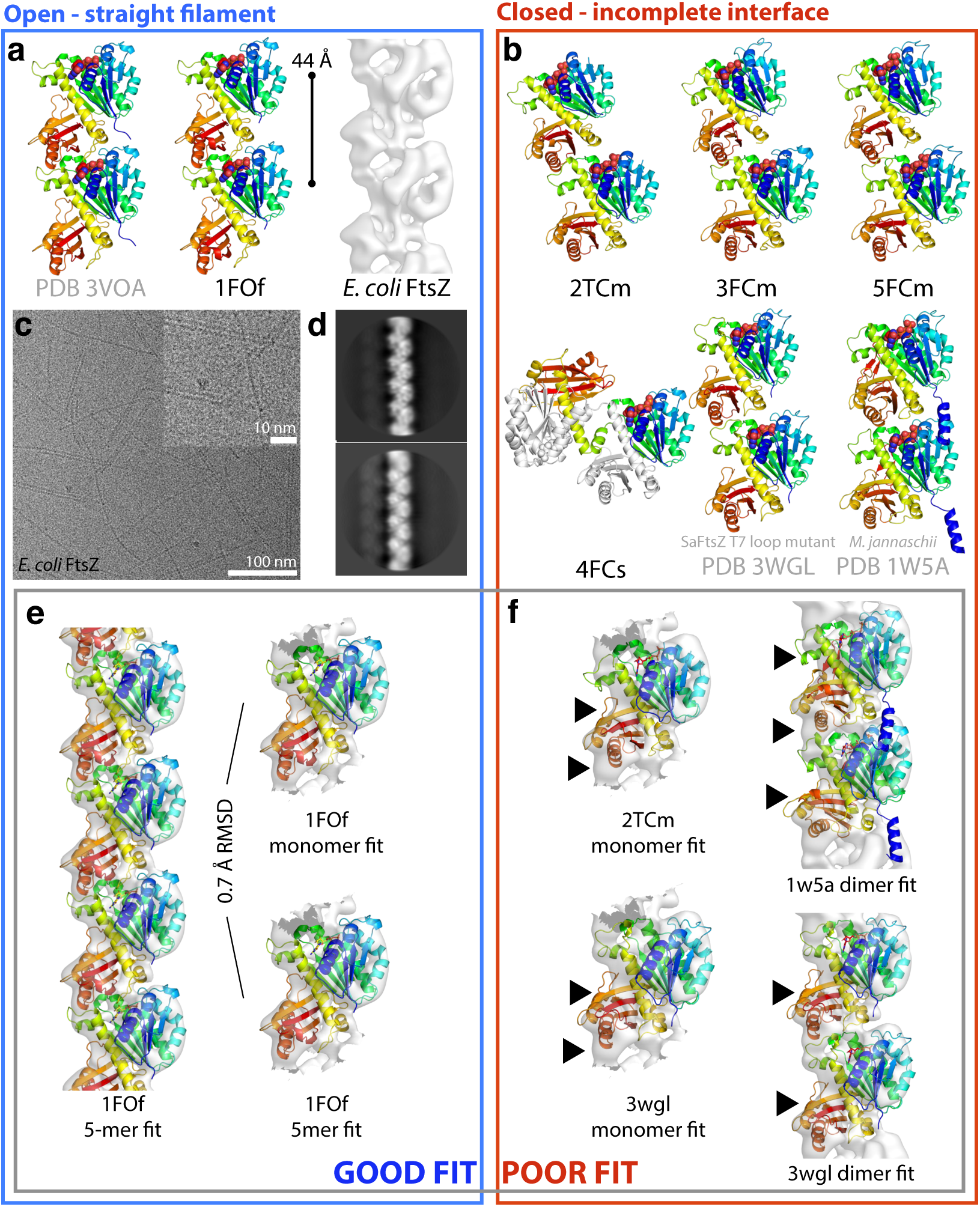
The closed conformation corresponds to the monomeric state of FtsZ, the open conformation corresponds to the filament, including in *E. coli*. **A) and B)** FtsZ pairs were extracted from crystal lattices as described in the text. Structures are shown in cartoon representation, each chain is rainbow coloured blue to red, N terminus to C terminus. Nucleotide atoms are coloured by element. In each case the view is from the same orientation after the lower molecule is aligned to the NTD of the lower subunit from PDB ID 3VOA. 4FCs is shown with one chain coloured and the other in white to highlight the domain swap. EcFtsZ filament cryoEM density is shown at a threshold of 7.5 σ, the same as in D. Open structures (panel A) can be arranged to have 44 Å repeats using the favourable tubulin-like interfaces found in crystals 3VOA and 1FOf. Closed structures (B) have smaller, incomplete, inter-subunit interfaces within crystals and cannot be sensibly arranged to produce straight filaments with a 44 Å repeat. **C)** Typical micrograph of frozen-hydrated EcFtsZ.GMPCPP filaments. Curved, straight, single and double/bundled filaments are seen. Inset shows magnified view. 44 Å repeat is visible by eye. **D)** Representative EcFtsz filament 2D classes produced by RELION **E, F)** FtsZ structures as indicated were fitted into the EcFtsZ cryoEM density using the CHIMERA volume viewer fitting tool. (E) A 1FOf 5-mer fits very well into the density, as does a 1FOf monomer – with both fits extremely similar. RMSD is for middle subunit in rigidly fitted 5-mer and monomer fitted into middle subunit density (F) Closed structures do not fit well into the electron density, and certainly not so that a repeating filament can be constructed. Some regions of poor fit are indicated with black arrowheads.

As discussed, previous work generated SaFtsZ structures in the closed form by extensive alteration of the T7 loop (PDB IDs 3WGL, 3WGK). These structures contain SaFtsZs that are clearly in the closed conformation (Figure 2D), but are arranged in straight filaments in the crystal. However, these filaments are not the same as the open form filaments, with a much smaller interface buried surface area of ∼700 Å^2^ as compared to ∼1200 Å^2^ for 1FOf and PDB ID 3VOA (calculated with PDBePISA server (35)), and a repeat of 45 Å. A dimer from a 3WGL pseudofilament is shown in Figure 5B. The longitudinal contact is made between residues from the bottom subunit at the N−terminus of H5 and the preceding loop (including residue F138), and the loop between H6 and H7. From the top subunit the T7 loop (replaced in these structures), one face of S9, and the loop at the N−terminus of H10 are involved. Interaction does not involve any of the residues on the other side of the interface, towards the phosphates of the nucleotide.

In contrast, we present four crystal forms where, for the first time, SaFtsZ is not arranged in straight, infinitely long filaments. In three of these, 2TCm, 3FCm and 5FCm we find in each case that one of the molecules in the ASU forms what looks superficially like a filament interface via its top face, and the other molecule in the ASU equivalently contributes a bottom face to another pseudo-interface. The crystals are therefore composed of pairs of poorly-interacting FtsZs (shown in Figure 5B) which pack via further crystal contacts that do not resemble interfaces. The pseudo-interfaces have subunit-subunit buried surface area (BSA) of 670-800 Å^2^, and look similar to the interfaces seen in the closed T7 mutant structures, only including residues from one side of the top face.

We know that our mutations inhibit filament formation (Figure 1), and we obtained crystals where these mutant SaFtsZs adopt the closed conformation, and fail to form *bona fide* interfaces. Hence we propose that these closed forms correspond to the conformation of monomeric SaFtsZ in solution. The pseudo-interface seen is perhaps best seen in this context as a consequence of crystallisation, which uses conditions that enhance protein-protein interactions, and not a cause. If the protein will crystallise it is extremely likely that one of the major crystal contacts will imitate the longitudinal interface, as the interface regions are most likely sticky. However, we cannot rule out the possibility that this minimal interface *(in silico* repetition of which generates a highly curved filament) is a functionally relevant and stable way for FtsZs to interact in solution.

The fact that the phosphate end of the interface is not formed in any of the F138A closed form crystals supports the idea that it is the closed conformation that is not compatible with formation of *bona fide* interfaces, not the mutation per se – because the F138A mutation is within the pseudo-interface. The fact that we recovered the 1FOf crystal form for the F138A mutant suggests that the conformational switch in this mutant is finely balanced.

The fourth closed form structure, 4FCs, as already mentioned, is arranged very differently within the crystal. The two pseudo-whole domain-swapped FtsZs formed by each pair of polypeptides both clearly adopt the closed conformation (Cα RMSD for comparable atoms in pseudo-monomer vs 2TCm 1.0 Å). The domain-swapped FtsZs do not make any crystal contacts that resemble filament interfaces. That a domain swap can happen, and that a domain-swapped FtsZ adopts a closed conformation, suggests that the two domains have a significant degree of independence and, more importantly here, that when SaFtsZ conformation is not modulated by polymerisation or pseudo-interfaces, the molecule adopts the closed conformation, as suggested previously by molecular dynamics (24).

We are able to significantly bolster the case that polymerisation into a straight filament is the driving force for the conformational switch we observe in crystal structures by presenting a medium (∼8 Å) resolution electron cryomicroscopy (cryoEM) structure of wildtype, full-length *E. coli* FtsZ straight filaments, Fig 4A,C,D. These data suffer from information anisotropy due to poor recovery of certain filament orientations in micrographs (Figure 5C). However, they clearly reveal an FtsZ filament with a 44 Å repeat and a density envelope into which SaFtsZ open conformation filaments can be fitted very satisfactorily, and closed form structures cannot be fitted (neither as pseudo-dimers nor monomers, *M. jannaschii* closed conformation dimer structure PDB ID 1W5A is shown fitted as it has previously been suggested to represent the conformation of FtsZ filaments (30)) (Figure 5D, and Videos S2 and S3). Several secondary structure elements can be unambiguously identified in the reconstruction, including H7 and, crucially, the planes of both N−and C-terminal domain beta sheets showing that the molecule is clearly in the open conformation (Video S2).

Given that all FtsZ crystals where *bona fide* straight filaments are seen in the crystal have FtsZs in the open conformation, and that the inverse is true – SaFtsZ filament crystals contain open subunits, and that our intermediate resolution EcFtsZ cryoEM structure also contains open subunits, we propose that a polymerisation-associated conformational switch is a general property of FtsZs. One of the consequences of such a switch, namely permitting cooperative assembly of a single-stranded filament, have been discussed previously (23, 26, 27, 36), however, such a switch confers additional surprising properties.

### The FtsZ conformational switch between monomer and filament provides filament-end asymmetry necessary for treadmilling

In preparing this manuscript it became clear to us that theoretical considerations of treadmilling can be fraught with intellectual traps. Having run the gauntlet of these potential pitfalls we present simplified, yet robust, schema to explain how a polymerisation-associated conformational switch provides the end-asymmetry necessary for treadmilling within a single stranded filament. We focus on the specific case where nucleotide forms part of the filament interface (i.e. in a tubulin-like fashion). In these, solvent exposed NDPs (nucleoside diphosphate) are quickly exchanged with NTPs (nucleoside triphosphate), NTP hydrolysis is not immediate, and NTP interfaces are stronger than NDP interfaces; although many of the conclusions are the same for filaments where nucleotide is buried inside subunits and has an allosteric effect on interface strength (i.e. actin-like).

Treadmilling of cytoskeletal filaments is a useful dynamic property. Treadmilling filaments can be used to push or pull molecules in the cell without motor proteins as long as end-tracking mechanisms or co-factors exist, and these behaviours can be made switchable with high flux through the filament (e.g. in eukaryotic anaphase microtubules). Recently it has been shown that FtsZ can treadmill *in vitro* with FtsA (16) and also alone (15), and that FtsA/Z treadmilling in cells guides septal cell wall remodelling (17, 18). FtsZ is widely considered to function as a single filament. However, as has been noted previously (19), a single-stranded filament with the above properties and rigid subunits, without conformational changes, cannot do robust treadmilling.

Such a hypothetical filament with rigid subunits is shown in Figure 6A. Note that the location (top/bottom) of nucleotide binding to the monomer is not important. This filament treadmills if a nucleotide gradient along the filament exists, and the kinetic plus end (net growth) will be at the end with more NTP. On-rates cannot differ at the two ends because they are the same reaction, but off-rate at the minus (GDP) end will be greater than at the plus end, so a situation of net growth at one end and net shrinkage at the other can be produced by changing monomer concentration (addition reaction is 1^st^ order with respect to monomer, loss is 0 order). This does not represent robust treadmilling however, as breakdown of terminal GDP interfaces is equivalent to breakdown of a GDP interface anywhere in the filament – and these processes will occur at the same rate as they are all 0 order. As noted previously (37) filament breakage and annealing could be an important facet of dynamics, but the filament in Figure 6A has a more fundamental limitation on it’s biological usefulness: the direction of treadmilling is determined entirely by the history of the filament (the direction of the initial NTP-NDP gradient), so there is no coupling of kinetic and structural polarity – and the same filament could treadmill in either structure-defined direction.

**Figure 6.**
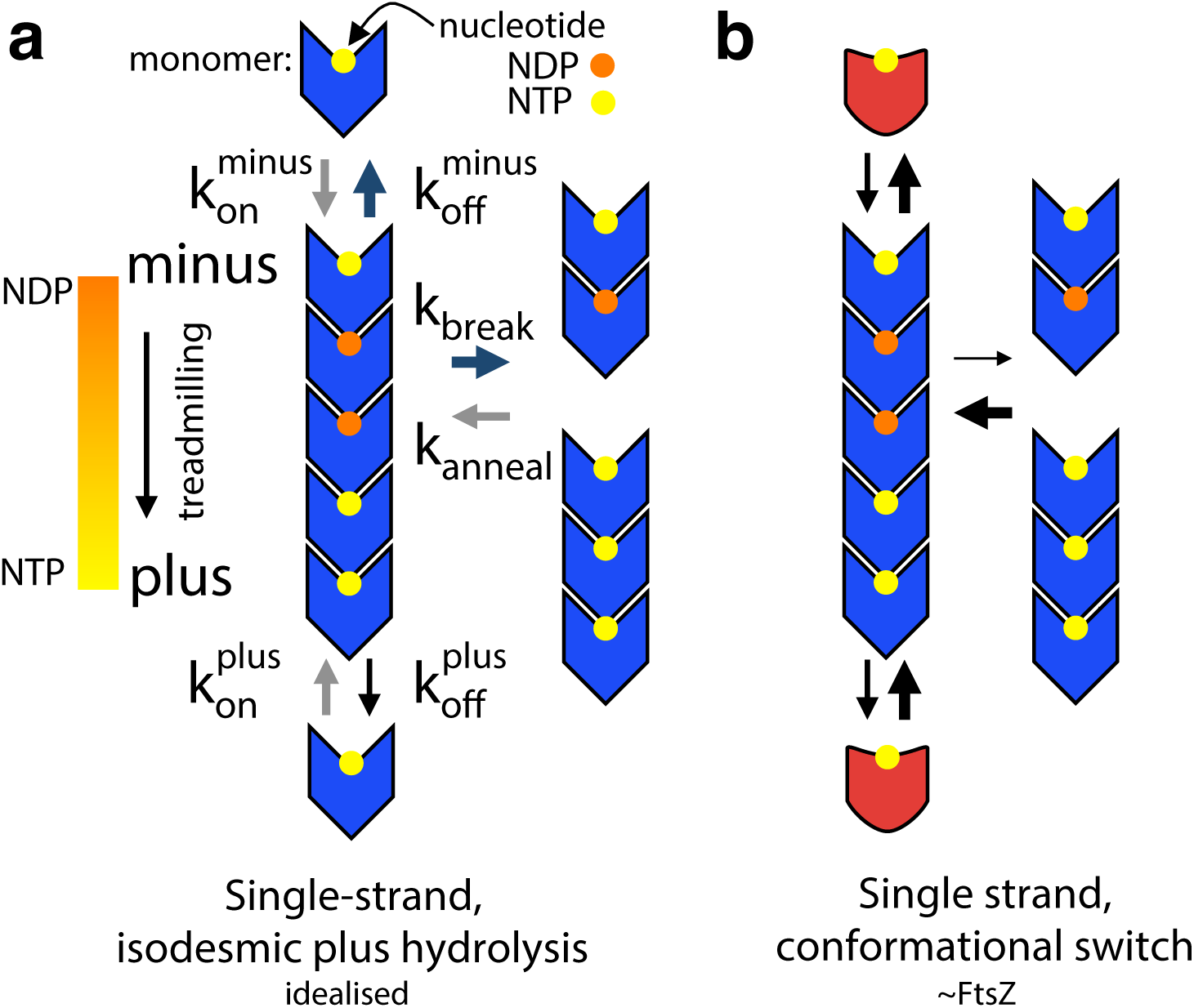
FtsZ’s polymerisation-associated conformational switch allows treadmilling of single-stranded filaments. Black arrows indicate rates roughly in proportion to their width, coloured arrows in (A) indicate rates that are equivalent. See main text for discussion of limitations and assumptions of these simplified models, particularly regarding implied orientation of molecules. **A)** An idealised rigid (lacking a conformational switch), tubulin-like, filament forming protein, for which addition/loss of a given NXP is isodesmic. This filament canot do robust treadmilling, as breakage is the same as minus end subunit loss, and it cannot couple structural and kinetic polarity **B)** A singlestranded version of (A) with a polymerisation-associated conformational switch, able to treadmill robustly and with coupled kinetic and structural polarities. The conformational switch allows filament breakage and subunit loss from ends to be different, and for the stereochemistry of subunit addition at either end to be different – meaning that addition will take place at different rates in a manner defined by structural polarity.

Coupling of kinetic and structural polarity requires subunit addition and or loss to proceed via different stereochemical pathways at each structure-defined end of the filament. This is not the case in Figure 6A, the difference in off-rates is defined by the nucleotide gradient and not structural polarity, and we have already seen that there can be no difference in on-rates. Filament systems can generate different stereochemistry for subunit addition at either end by being multi-stranded and having staggered subunits that undergo a conformational change, such as actin (38), or by using a longitudinal hooking mechanism, such as TubZ (39).

Figure 6B shows our model for how a single filament very similar to the case in Figure 6A can couple its structural polarity to a defined kinetic polarity and thus usefully, and robustly, treadmill. The crucial difference between Figure 6A and B is the existence of a polymerisation-associated conformational switch, i.e. subunits are no longer rigid, but can exist in one of two conformations – one form associated with the polymer, the other adopted in the free monomer. The free energy cost of the conformational switch from closed to open is paid for by binding to a filament end, and in the other direction through nucleotide hydrolysis and exchange that makes the longitudinal NDP intersubunit interface unfavourable. Although formation of a NTP interface at either structurally-defined end has the same net energy change, the reaction pathways are stereochemically different, and will occur at different rates because two different pairs of molecular surfaces are involved in each case initially. This difference is illustrated in a non-rigorous fashion in the context of our structures in Figure 7.

**Figure 7.**
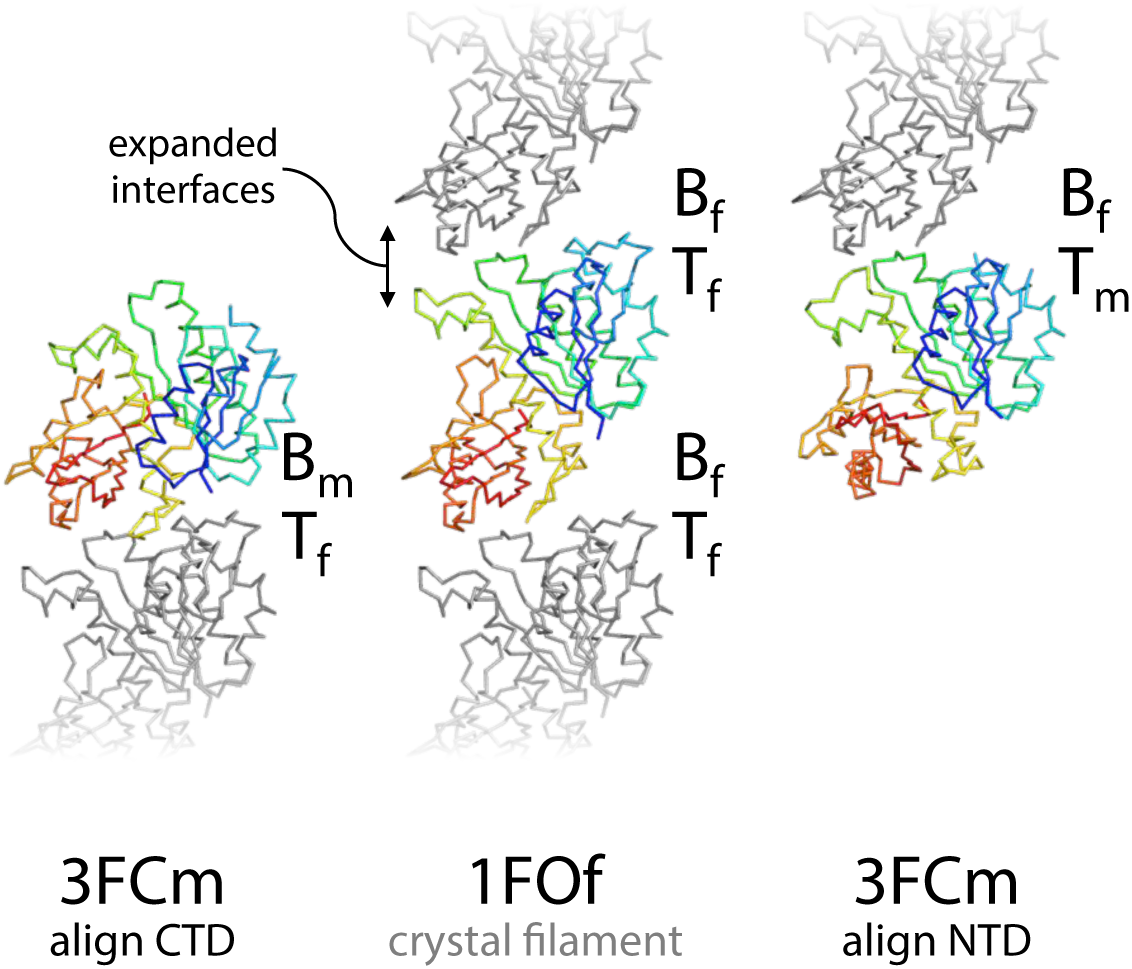
A polymerisation-associated conformational switch generates chemical asymmetry at filament end interfaces. (Middle) 3 molecules from the open form 1FOf crystal filament, slightly separated for clarity, are shown as Cα ribbons. The middle subunit is rainbow coloured blue-red, N− to C-terminus, the top and bottom subunits are coloured grey. (Right, left) The middle subunit is replaced with a closed form 3FCm molecule, aligned to the middle subunit N−terminal domain (NTD) (right) or C-terminal domain (CTD) (left), indicated by the arrows. The different pairs of approaching surfaces are labelled B/T_m/f_: bottom/top, monomer/filament. These modelled closed-open interfaces will not represent the transition state of subunit addition at either end of a filament (nor even any position on the reaction pathway), but they illustrate the fact that the conformational switch will necessarily lead to stereochemically different reaction pathways at each end that allow the two ends to have different rates of subunit addition, linking structural and kinetic polarity.

Importantly, the scheme in Figure 6B also allows breakage at NDP interfaces within the filament to be different to loss of NDP subunits from each end: essentially NDP interfaces in the filament can be stronger because the energetic cost of breaking them, when losing subunits from ends only, is paid for by the favourable switch to the monomer conformation which the end subunits of two halves of the broken filament cannot adopt because they remain in filaments. Especially important to note here that the scheme in Figure 6B can also be drawn with nucleotide on the other side of the monomer, i.e. we are not making a prediction about which end of a single-stranded FtsZ filament is the kinetic plus end – an issue that will need further investigation (37, 40, 41).

## Conclusion

Here we have shown that FtsZs adopt two different conformations: open and closed. The open form is adopted by FtsZ in straight filaments, the closed form by FtsZ monomers. The implication is that the polymerisation-associated switch from closed to open is made favourable by the free energy gain of interface formation. Such a polymerisation-associated conformational switch explains how a single stranded filament can show cooperativity in polymerisation, and how a single-stranded filament can avoid breaking apart when treadmilling. This switch also explains how a single-stranded filament with tubulin-like properties can couple structural and kinetic polarities to enable useful treadmilling.

At this point it should be noted that although single-stranded FtsZ is frequently considered the functional unit of the protein *in vivo*, the evidence for this could be stronger. Nevertheless, the potential of the conformational switch to generate end-asymmetry could also be exploited in multistranded treadmilling. In addition to this, we have not directly addressed the structural basis of filament bending. Indirectly, cryoEM of *E. coli* FtsZ filaments assembled with GMPCPP and frozen after a 20-30 second incubation show some degree of bending in almost all single filaments, and segments from bent filaments are included in the reconstruction showing subunits in the open form — apparently undermining previous ideas that all bent FtsZ filaments are GDP-bound and/or in the closed conformation.

## Materials and methods

### Cloning, protein expression and purification of FtsZ

Full-length (FL) *SaFtsZ* (Uniprot ID: FTSZ_STAAU) was amplified using PCR from genomic DNA and cloned into the NdeI and SapI sites of pTXB1 (NEB IMPACT system, NEB Ipswich, MA), thus generating a C-terminal fusion to the *Mxe* intein/chitin binding domain which self-cleaves upon the addition of DTT. PCR mutagenesis using this vector as a template generated mutants T66W and F138A. Full-length fusion proteins were expressed in *E. coli* BL21 (DE3) cells, which were grown in LB media with ampicillin (100 mg/L) at 37 °C with 200 rpm shaking to an OD_600_ of 0.6. Cultures were then shifted to 16 °C and after 1 hour expression were induced by the addition of 0.4 mM isopropyl-β-D-1-thiogalactopyranoside (IPTG), before overnight incubation. Cells were collected by centrifugation and resuspended in buffer FL.A (50 mM HEPES-KOH, 50 mM NaCl, 20mM EDTA, pH 8.5) with 100 μg/mL lysozyme, 10 μg/mL DNAse and 4 mg/mL PMSF, before lysis via 2-3 passes through a French press. Lysate was clarified by centrifugation at 100,000 g and 4 °C for 1 hour. Soluble protein was captured on a chitin column (NEB) equilibrated and washed with buffer FL.A. Intein activity and release of the untagged, full-length protein was initiated by overnight incubation in buffer FL.B (buffer FL.A with 50 mM DTT) at 4 °C followed by elution. Eluate was further purified by anion-exchange chromatography on a 5 mL HiTrap Q column (GE Healthcare, Little Chalfont, UK). The column was equilibrated and washed with buffer FL.Q.A (50 mM Tris-HCl, 1 mM EDTA, pH 7.5) and bound protein was eluted with a linear gradient of buffer FL.Q.B (FL.Q.A with 1 M NaCl). Peak fractions were further purified by gel filtration on a 70 mL Sephadex 75 (GE Healthcare) column in Buffer FL.GF (20 mM Tris-HCl, 150 mM NaCl, 10% glycerol, pH 8.0). Peak fractions were pooled and concentrated using centrifugal concentrators (Vivaspin, Sartorius, Epsom, UK) to 5-8 mg/mL before freezing in liquid nitrogen and storage at −80 °C. Protein integrity was confirmed by electrospray masspectrometry.

Truncated (TR) (12-316), *SaFtsZ* (Uniprot ID: FTSZ_STAAU) was cloned into a pHis17 plasmid derivative, with no tag, via Gibson assembly techniques. PCR mutagenesis using this vector as a template generated mutants T66W and F138A. Truncated *SaFtsZ* proteins were expressed in *E. coli* C41 (DE3) cells (Lucigen) grown in 2xTY media with ampicillin (100 mg/L) at 37 °C with 200 rpm shaking to an OD600 of 0.6. Cultures were then shifted to 16 °C and after 1 hour expression was induced by the addition of 0.5 mM IPTG, before overnight incubation. Cells were collected by centrifugation and resuspended in Buffer TR.A (50 mM Tris-HCl, 30 mM NaCl, pH 8.0) before lysis by passing through a cell disruptor (Constant Systems, Daventry, UK) at 25 kpsi. The lysate was clarified by centrifugation at 100,000 g and 4 °C for 30 minutes. The soluble fraction was loaded onto a HiTrap Q anion-exchange column (GE Healthcare), washed with Buffer TR.Q.A, and eluted with a linear gradient of Buffer TR.B (TR.A with 1 M NaCl). Peak fractions were pooled, and protein was precipitated by adding saturated ammonium sulphate to 35% v/v. Precipitated protein was centrifuged at 28,000 g and 4 °C for 30 minutes, and the pellet resuspended in Buffer TR.A. Resuspended protein was further purified by size-exclusion chromatography on a HiLoad Sephacryl S300 16/60 column (GE Healthcare) in Buffer TR.A. Peak fractions were pooled, concentrated to 15-25 mg/mL using centrifugal concentrators (Vivaspin, Sartorius) before freezing in liquid nitrogen and storage at −80°C. Protein integrity was confirmed by electrospray masspectrometry.

Full length, untagged, *E. coli* FtsZ (Uniprot ID: FTSZ_ECOLI) was cloned into the BamH/NdeI sites of pET9a (Novagen), with no tag. Purification of *E. coli* FtsZ was by a modified version of established protocols (42). Protein was expressed in *E. coli* C41 (DE3) cells (Lucigen) grown in 2xTY media with kanamycin (50 mg/L) at 37 °C with 200 rpm shaking to an OD_600_ of 0.6. Cultures were then shifted to 20 °C and after 1 hour expression was induced by the addition of 0.5 mM IPTG, before overnight incubation. Cells were collected by centrifugation and resuspended in buffer PEM (50 mM PIPES-KOH, 5 mM MgCl_2_, 1 mM EDTA, pH 6.5) before lysis by passing through a cell disruptor (Constant Systems) at 25 kpsi. The lysate was clarified by centrifugation at 100,000 g and 4 °C for 30 minutes. FtsZ was separated by GTP and calcium-induced precipitation. Lysate was adjusted to pH 7 with acetic acid then GTP and CaCl2 were added to 1 mM and 20 mM respectively. This mixture was then centrifuged at 11,000 g and 4 °C for 15 minutes. The pellet, containing FtsZ, was resuspended in buffer PEM and insoluble debris was removed by centrifugation at 100,000 g and 4 °C for 30 minutes. FtsZ was further purified by anion exchange chromatography over a Mono Q 4.6/100 (1.7 mL) PE column (GE Healthcare) equilibrated and washed with buffer ECZ.Q.A (50 mM Tris-HCl, 50 mM KCl, 1 mM EDTA, 10% glycerol, pH 8.0), bound protein was eluted with a linear gradient of buffer ECZ.Q.B (Buffer ECZ.Q.A with 1 M KCl). Peak fractions were pooled, concentrated to 20 mg/mL using centrifugal concentrators (Vivaspin, Sartorius) before freezing in liquid nitrogen and storage at −80°C. Protein integrity was confirmed by electrospray masspectrometry.

### Sedimentation analysis of assembly

Samples were prepared in a Thermostat Plus (Eppendorf, Hamburg, Germany) at 25 °C, where the protein (10 μM) was equilibrated in Buffer HKM (50 mM HEPES-KOH, 100 mM potassium acetate, 5 mM magnesium acetate, pH 7.7). After the addition of 1 mM GTP or 0.1 mM GMPCPP, with or without 20 μM PC190723 and 2% DMSO to prevent ligand precipitation, samples were incubated for 20 minutes at 25 °C and then centrifuged at 400,000 x g for 20 minutes at the same temperature in a Beckman TLA 100 rotor. The supernatants were carefully withdrawn, and the pellets were resuspended in the same volume of buffer. Subsequently, pellets and their corresponding supernatants were loaded and run in the same well of an SDS-PAGE gel with a 25 minutes delay to analyse the amount of protein polymerised in the pellet versus the unassembled fraction in the supernatant.

### GTPase activity assay

GTP (Sigma, St Louis, MI), and guanosine-5’-[(α,β)-methyleno]triphosphate (GMPCPP) (Jena Bioscience, Jena, Germany) hydrolysis by full-length *SaFtsZ* proteins was monitored by detecting the release of inorganic phosphate with the malachite green assay (43) in samples containing 10 μM or 20 μM of protein in Buffer HKM at 25 °C. When included, PC190723 was at a concentration of 20μM and samples contained 2% DMSO to prevent ligand precipitation

### Negative stain electron microscopy

Full-length wild-type and mutant *SaFtsZ*s were visualised by negative-stain electron microscopy. About 20 μL of sample, prepared as for sedimentation analysis of assembly (except protein was at 20 μM), were applied to formvar and carbon-coated copper grids, incubated for one minute, and then stained with 2% w/v uranyl acetate in water. Images were taken at several magnifications using a JEOL 1200 EX-II microscope operated at 100 kV equipped with a Gatan CCD camera.

### Crystallisation

Crystallisation conditions were found using our in house high-throughput crystallisation platform, by mixing 100 nL truncated *SaFtsZ* T66W or F138A solution at 5 or 10 mg/mL, with GTP at 10 mM, with 100 nL of 1920 different crystallisation reagents in MRC vapour diffusion sitting drop crystallisation plates. Conditions yielding crystals were optimised, and crystals from either the initial screens or subsequent optimization were selected for data collection. Conditions giving the crystals for which structures are presented are in Table S1.

### Crystallographic data collection and structure determination

Diffraction images were collected from single frozen crystals at beamlines at either DLS (Diamond Light Source, Harwell, UK) or ESRF (European Synchrotron Radiation Facility, Grenoble, France) as indicated in Table S1, at 100 K. Diffraction images were processed with XDS, POINTLESS and SCALA software. Initial phases were determined by molecular replacement using PHASER with search models as indicated in Table S1. Models were rebuilt manually using MAIN and refined using REFMAC and PHENIX.REFINE. Ramachandran plots and MOLPROBITY statistics were used to validate the structures.

### Structure of E. coli FtsZ filament by cryoEM

For cryoEM, *E. coli* FtsZ was prepared at 0.5 mg/mL in 50 mM HEPES-KOH, 100 mM potassium acetate, 5 mM magnesium acetate, pH 7.7, at 20 °C. GMPCPP was added to 0.1 mM. 2.5 μL sample was applied to freshly glow-discharged Quantifoil Cu R2/2 200 mesh grid and plunge frozen using a Vitrobot Mark III (FEI Company, OR) into liquid ethane maintained at 93.0 K using an ethane cryostat (44). The Vitrobot chamber temperature was set to 10 °C and humidity to 100 %. Micrograph movies of FtsZ filaments were collected with an FEI Tecnai G2 Polara microscope operating at 300 kV. Data were acquired on a Falcon III direct electron detector protoype at a calibrated pixel size of 1.34 Å and an approximate total dose of 40 e^-^/Å^2^ using the automated acquisition software EPU (FEI Company). A total of 1834 movies were collected at −1 to −4 μm defocus in 46 frames during each 1.5 s exposure. All image processing was carried out within RELION 2.0 (45). Micrograph movies were motion corrected using MotionCor2 (46) with 5 x 5 patches and a grouping of 10 frames. Helical autopicking in RELION was used in order to find segments along the filaments at 4.3 nm intervals with confidence. Boxes of 190 x 190 pixels were extracted around each segment. After 2D classification, a 3D autorefinement of the remaining 511,000 filament segments was performed in RELION using helical reconstruction options, against an atomic protofilament model derived from PDB ID 3VO8, filtered to 20 Å. The resulting two half maps were used in post processing to sharpen the map (B-factor −360 Å^-1^) and to obtain a gold standard FSC-based resolution estimate of 7.8 Å (0.143 FSC criterion). In the absence of an *E. coli* FtsZ crystal structure, CHIMERA was used to determine that the SaFtsZ filament structure, showing the open conformation, fits very well into the E. coli filament density, as opposed to any other structure containing closed conformations.

### Structural figures

All structural figures were prepared in PyMOL (Schrödinger, Inc.), with some density volume operations carried out in CHIMERA (47).

## Funding Information

This work was funded by the Medical Research Council (U105184326 to JL), the Wellcome Trust (095514/Z/11/Z to JL) the Ramón y Cajal Program (RYC-2011-07900 to MAO) and MINECO (BFU2014-51823-R to JMA). JMW is the recipient of a Boehringer Ingelheim Fonds PhD Fellowship.

## Acknowledgements

We would like to thank Minmin Yu for assistance with data collection and Piotr Szwedziak for the *E. coli* FtsZ plasmid (both LMB).

## Author Contributions

MAO and AGS performed biochemical assays. JMW, MT, DKC and JL performed crystal structure determinations. JMW carried out cryoEM data collection and processing. JMW, MAO, JMA and JL analysed data and wrote the manuscript.

## Supplementary Material

**Table S1 – Crystallographic and cryoEM data**

**Video S1 – The SaFtsZ conformational switch** This video shows a morph interpolation between SaFtsZ structures 1FOf and 3FCm, from two angles, with and without side chains.

**Video S2 – EcFtsZ filaments contain subunits in the open conformation** A SaFtsZ 1FOf filament is shown fitted into the EcFtsZ cryoEM density at 7 sigma contour. A 1FOf monomer is shown fitted into the EcFtsZ density at 7,8 and 9 sigma, with two additional views demonstrating the unambiguous location of both N−and C-terminal domain beta sheets.

**Video S3 – Closed form monomers do not fit in the EcFtsZ filament density.** The 1FOf open monomer and the 2TCm closed monomer are compared after fitting into the cryoEM density. The poor fit of the 2TCm C-terminal domain is demonstrated further by comparing simulated EM density (prepared using the EMAN2 pdb2mrc utility, Tang, G et al. (2007), J Struct Biol., 157(1), pp. 38-46.)

**Table S1.**
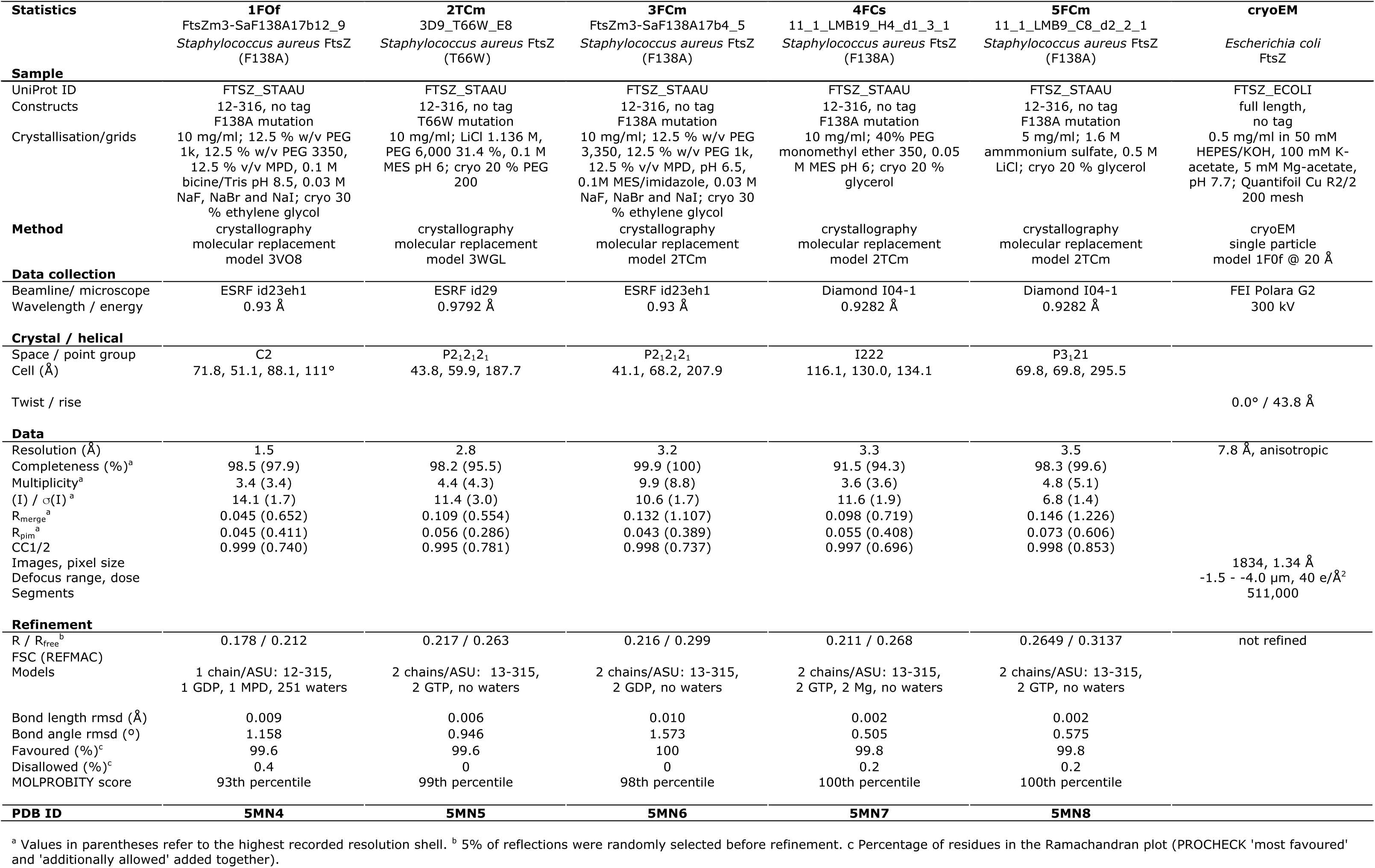
Crystallographic and cryoEM data

